# Fungal community composition along a gradient of permafrost thaw

**DOI:** 10.1101/2021.07.01.450738

**Authors:** Mariana Kluge, Christian Wurzbacher, Maxime Wauthy, Karina Engelbrecht Clemmensen, Jeffrey Hawkes, Karolina Einarsdottir, Jan Stenlid, Sari Peura

## Abstract

Thermokarst activity at permafrost sites releases considerable amount of ancient carbon to the atmosphere. A large part of this carbon is released via thermokarst ponds, and fungi could be an important organismal group enabling its recycling. However, our knowledge about aquatic fungi growing in thermokarstic systems is extremely limited. In this study, we collected samples from five permafrost sites distributed across circumpolar Arctic and representing a gradient of permafrost integrity. Samples were taken from the ponds surface water, the detritus and the sediment at the bottom of the ponds. These samples were extracted for total DNA, which was then amplified using primers targeting the fungal ITS2 region of the ribosomal genes. These amplicons were sequenced using PacBio technology. Surface water samples were also collected to analyze the chemical conditions in the ponds, including nutrient status and the quality and quantity of dissolved organic carbon. This dataset gives a unique overview of the impact of the thawing permafrost on fungal communities and their potential role on carbon recycling.

## Background & Summary

Frozen tundra soils hold one of the Earth’s largest pools of organic carbon. With ongoing climate change, permafrost is thawing rapidly, especially in the Arctic and Subarctic regions, causing the release of a large fraction of this carbon^1,2^. The thawing of permafrost creates small and shallow freshwaters, hereafter referred as thermokarst ponds^3^. The vast amount of organic matter released from degrading permafrost ends up in these ponds^4^, where it can sink and be stored in the sediment, or be recycled in the microbial loop, generating greenhouse gases (GHG) as end products^5,6^. Most of the research on the microbial activity in thermokarst ponds concentrates on prokaryotes^7–10^, and despite the central role of fungi as decomposers of the organic matter in terrestrial ecosystems^11–13^, very little is known about the fungal communities in thermokarst ponds. To our knowledge only one study has specifically targeted the fungi in thermokarst ponds, highlighting that a major part of aquatic fungal community growing inside sporadic permafrost region belongs to unknown phyla^14^.

In this dataset, we collected surface water, detritus and sediment from thermokarst ponds in five different permafrost areas in the Arctic, representing a gradient of permafrost degradation stages from unaffected permafrost sites (represented by Alaska and Greenland, hereafter referred as pristine) to sites affected by increasing severity of thermokarst activity (represented by Canada, Sweden and Russia, hereafter referred as degraded) (Figure 1). For each site, 12 ponds were sampled. Moreover, in the Canadian site, the ponds represented three different stages of permafrost thaw, including emerging, developing and mature thermokarst ponds; four ponds were sampled at each of these three stages^8,14^. This allowed us to investigate whether there is a succession of the community over pond development, while the quality and availability of carbon sources change. All water samples were extracted for the total metagenomic DNA, which was then amplified for fungal ITS2 region of the ribosomal genes and sequenced using PacBio (Figure 2). We also collected water samples for chemical and optical analyses, in order to investigate nutrients and GHG concentrations as well as the quantity and quality of the dissolved organic matter (DOM). This included nutrients (dissolved nitrogen (DN), nitrate (NO_3_^−^), nitrite (NO_2_^−^), ammonium (NH_4_^+^), sulfate (SO_4_^2−^), total phosphorous (total P)), total iron (Fe), GHG (carbon dioxide (CO_2_), methane (CH_4_)) and dissolved organic carbon (DOC) concentrations, as well as various proxies of DOM such as fluorescence index (FI), freshness index (BIX), humification index (HIX), specific ultraviolet absorbance (SUVA_254_), spectral slope for the intervals 279–299 nm (S_289_) and average H/C and O/C.

**Figure 1.**
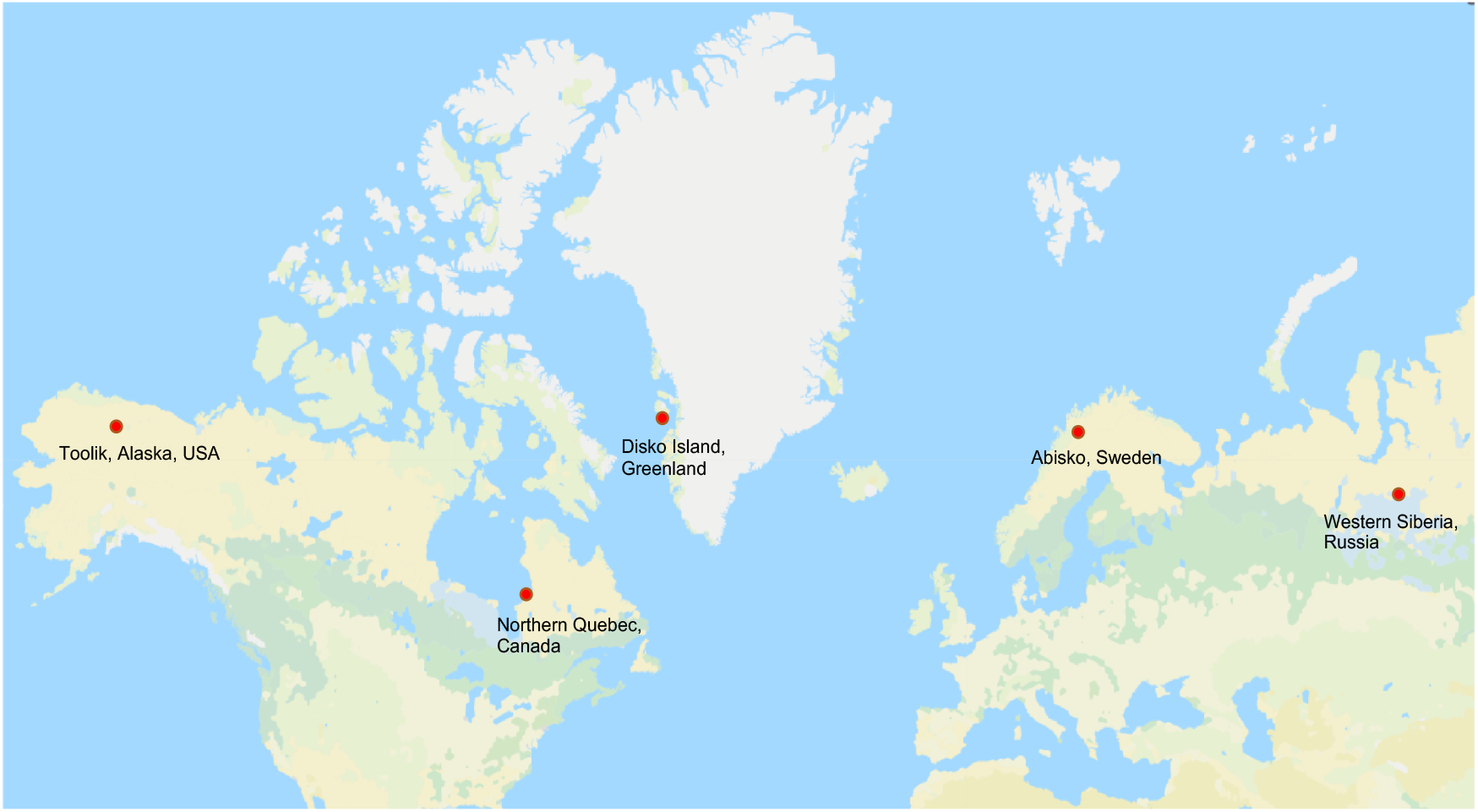
Locations of the sampling sites.

**Figure 2.**
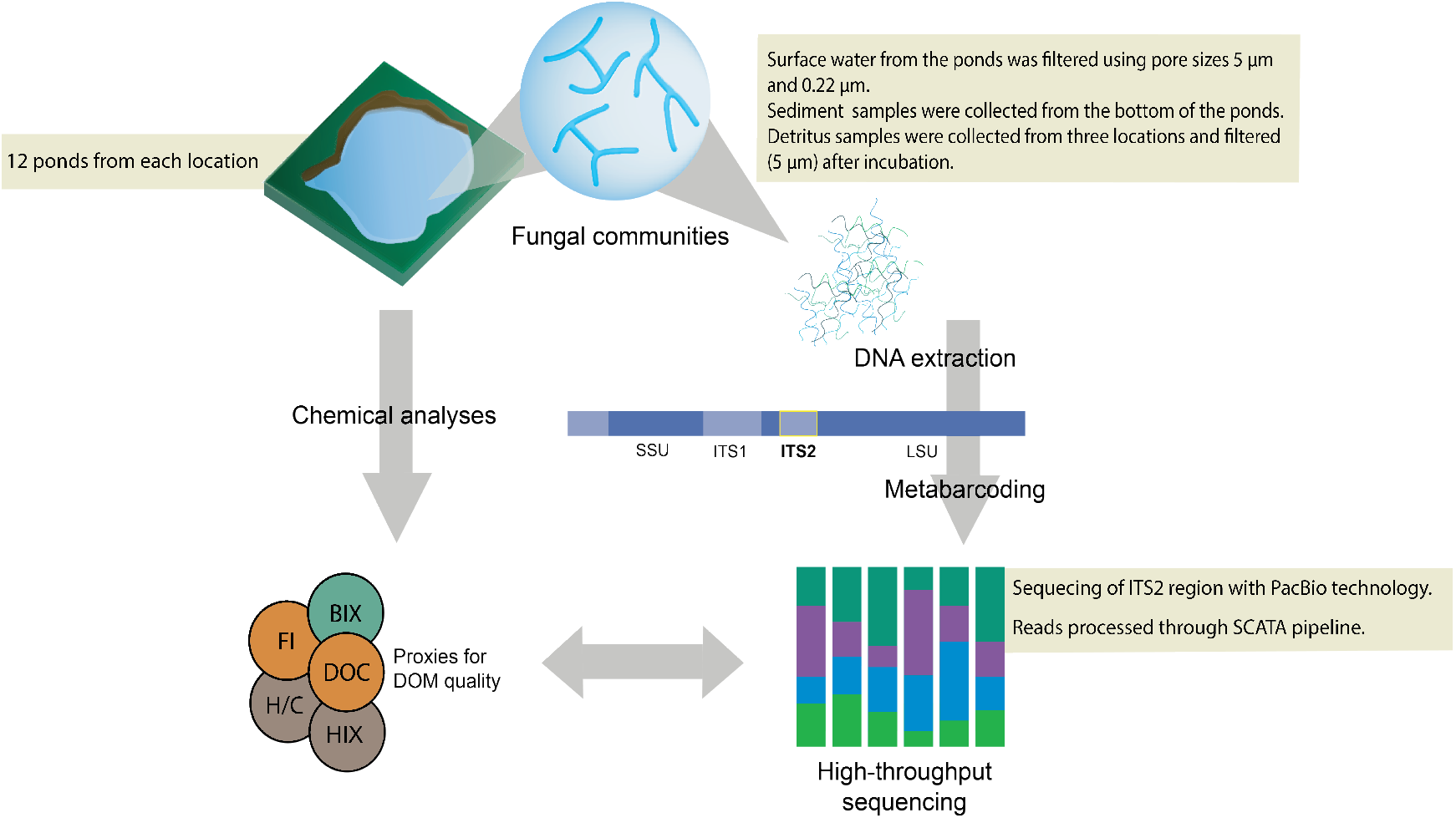
Workflow for producing the amplicon and chemical data.

The aim of this data collection was to study how the fungal communities are affected by permafrost thaw and resulting inputs of organic matter to the thermokarst ponds. Furthermore, the focus was also set on how the chemical conditions in the ponds may be linked to the fungal community composition. This dataset gives unique insight into the composition of fungal communities in aquatic habitats at sites representing different stages of permafrost integrity. The data can be used to study the general composition of arctic fungal communities and how the community changes together with their environment, such as availability of the carbon substrates. Also, it expands the database for fungal ITS sequences with a large number of previously unencountered sequences, widening the knowledge and database available for studying fungal diversity in undersampled biomes.

## Methods

### Study sites

We sampled a gradient of ponds in the following five sites representing different regional-scale permafrost integrity: Toolik, Alaska, USA; Qeqertarsuaq, Disko Island, Greenland, Denmark; Whapmagoostui-Kuujjuarapik, Nunavik, Quebec, Canada; Abisko, Sweden and Khanymey, Western Siberia, Russia (Table 1). The aim was to include representatives of different stages of permafrost thaw in order to understand whether responses can be generalized across different geographic and environmental conditions.

**Table 1.**
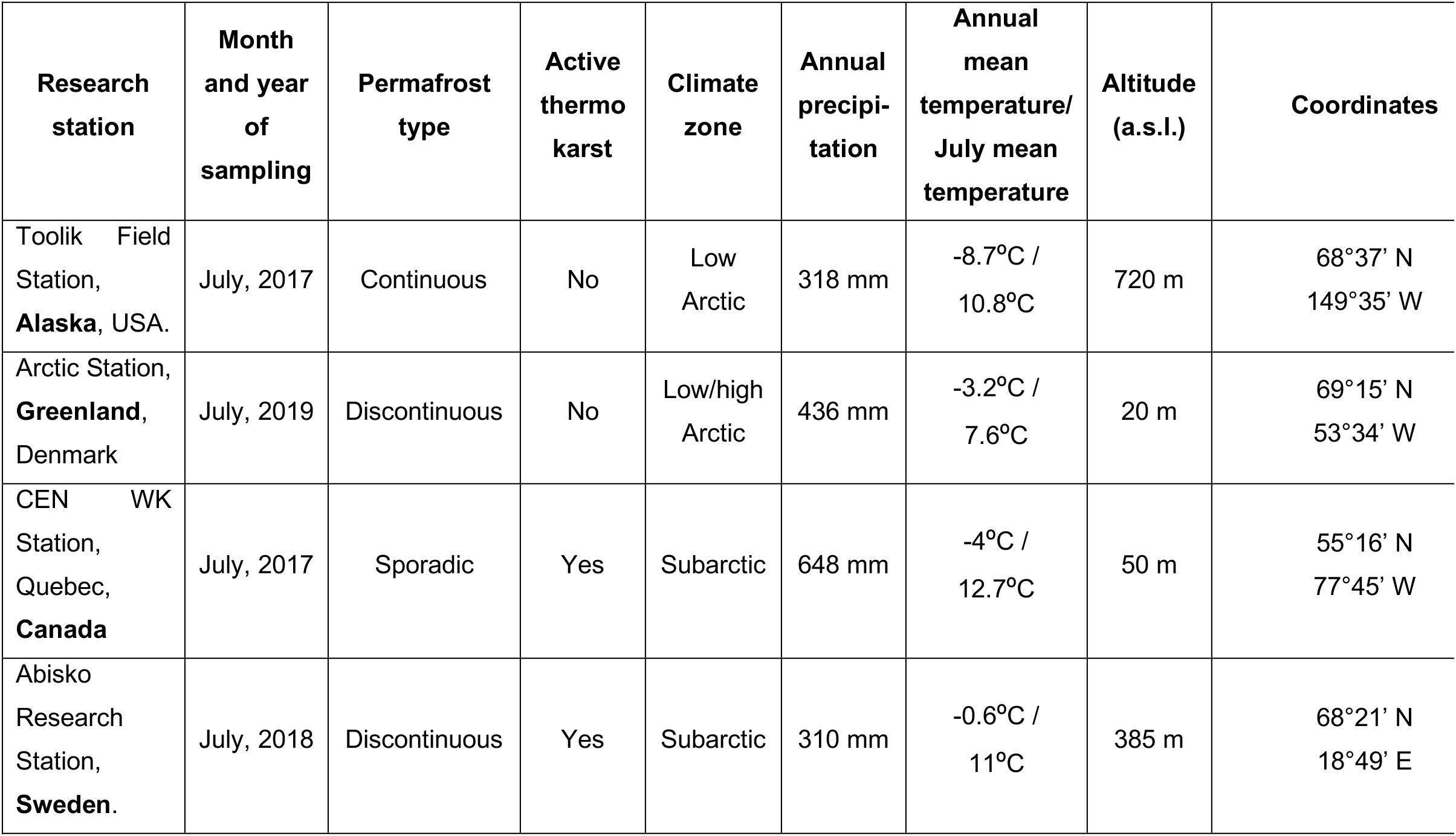
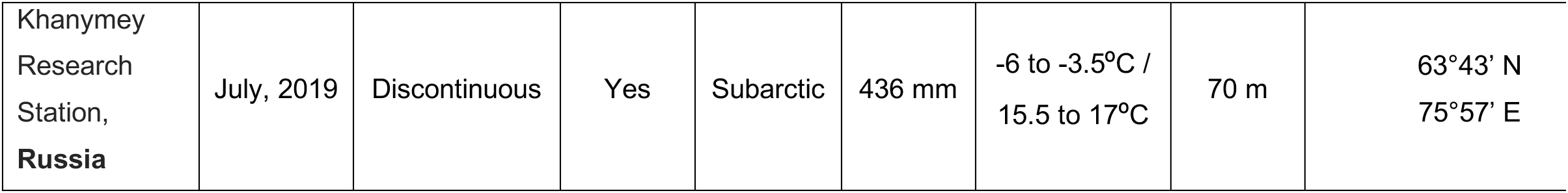
Description of the sampled sites. The stations are ordered according to increasing level of thermokarst activity. The coordinates are for the stations, the sample coordinates can be found at https://osf.io/hvwpu/?view_only=6f6b73e7522741cc9e350b0d25889db5. WK = Whapmagoostui-Kuujjuarapik Research Complex.

The sampling site in Alaska is located in a continuous permafrost area, mostly dominated by moss-tundra characterized by tussock-sedge *Eriophorum vaginatum* and *Carex bigelowii*, and dwarf-shrub *Betula nana* and *Salix pulchra*^15^. The average depth of the active layer for 2017 was ~50 cm^16^. Records of surface air temperature from 1989 to 2014 showed no significant warming trend, neither a significant increase of the active layer mean maximum thickness or maximum thaw depths^17^.

The sampling site in Greenland is located in the Blæsedalen Valley, south of Disko Island, and is characterized as a discontinuous permafrost area. From 1991 to 2011, Hollensen et al. (2015)^18^ observed an increase of the mean annual air temperatures of 0.2 °C per year in the area, while Hansen et al. (2006)^19^ highlighted a reduction of 50% of sea ice cover from 1991 to 2004. Soil temperatures recorded by the Arctic Station from the active layer of the coarse marine stratified sediments also show an increase over the years^18^. The sampling site is comprised of wet sedge tundra, and the dominating species are *Carex rariflora*, *Carex aquatilis*, *Eriophorum angustifolium*, *Equisetum arvense*, *Salix arctophila*, *Tomentypnum nitens* and *Aulacomnium turgidum* ^20^.

The Canadian site is located in a river valley within a sporadic permafrost zone, in a palsa bog. The vegetation consists of a coastal forest tundra, dominated by the species *Carex* sp. and *Sphagnum* sp. ^21^. Since the mid-1990s, there has been a significant increase in the surface air temperatures of the region for spring and fall, which has been correlated to a decline of sea ice coverage in Hudson Bay^22^. This area has experienced an accelerated thawing of the permafrost over the past decades, resulting in the collapsing of palsa and the emergence of thermokarst ponds and significant peat accumulation^21,23^. In this specific site, thermokarst ponds at different development stage can be found, from recently emerging new to older, mature thermokarstic waterbodies. The stage of the pond was estimated based on the distance between the pond and the edge of the palsa, as well as satellite pictures^14^. Emerging ponds have a maximum distance from the palsa of 1 m and were less than 0.5 m deep, whereas developing ponds have a distance from 2-3 m and were ~1 m deep. Mature ponds were identified based on satellite images and were up to 60 years old.

The Swedish site is located on a discontinuous permafrost zone at the Stordalen palsa mire, in on an area of collapsed peatland affected by active thermokarst. The region has experienced an increase in air annual mean temperatures and active layer thickness since the 1980s, which also was followed by a shift to wetter conditions^24^. The vegetation found on the palsa depressions of Stordalen mire is dominated by sedges (*Eriophorum vaginatum*, *Carex* sp.) and mosses (*Sphagnum* sp.) ^24,25^.

The Russian site is located in a discontinuous permafrost area in Western Siberia Lowland, near Khanymey village. The sampling site is a flat frozen palsa bog with a peat depth no deeper than 2 m, and is affected by active thermokarst, resulting in the emergence of thermokarst ponds^26,27^. The vegetation is dominated by lichens (*Cladonia* sp.), schrubs (*Ledum palustre*, *Betula nana*, *Vaccinium vitis-idaea*, *Andromeda polifolia*, *Rubus chamaemorus*) and mosses (*Sphagnum* sp.) ^28^.

### Sample collection

A total of 12 ponds were sampled per sampling site (60 ponds in total). We collected water samples from the surface to measure total P. Surface water was also filtered through GF/F glass fiber filters (0.7 μm, 47 mm, Whatman plc, Maidstone, United Kingdom) in order to analyze Fe, various dissolved anions and cations, and DOC concentrations, and perform optical and mass spectrometry analyses on DOM. Moreover, water samples were collected in order to measure GHG (CO_2_ and CH_4_) concentrations. Surface water, detritus and sediment samples were also collected from ponds for fungal community analyses. Water samples were collected and filtered first through 5 µm Durapore membrane filter (Millipore, Burlington, Massachusetts, USA) and then through a 0.22 µm Sterivex filter (Millipore) to capture fungal cells of different sizes. The samples were filtered until clogging or up to a maximum of 3.5 liters (filtered volume ranging from 0.1 l to 3.5 l). Sediments were sampled from the bottom of each pond, with the exception of Canadian site, where only one emerging and three developing ponds were sampled for sediment. From the sites in Greenland, Sweden and Russia, detritus samples (dead plant material) were taken. The detritus was washed in the lab using tap water, followed by overnight incubation in 50 ml tap water to induce sporulation. After the incubation, the water was filtered through a 5 μm pore size filter.

The DNA samples were transported to the laboratory frozen, with the exception of the Alaskan samples, which were freeze dried prior to transportation. The samples transported frozen were freeze dried prior to DNA extraction to ensure similar treatment of all samples. The samples for nutrient and carbon measurements were transported frozen with the exception of samples for DOC and fluorescence analyses, which were transported cooled.

### Chemical analyses

All chemical, optical and mass spectrometry results are provided in https://osf.io/hvwpu/?view_only=6f6b73e7522741cc9e350b0d25889db5. DOC quantification was carried out using a carbon analyzer (TOC-L + TNM-L, Shimadzu, Kyoto, Japan). Accuracy was assessed using EDTA at 11.6 mg C/l as a quality control (results were within +− 5%) and the standard calibration range was of 2-50 mg C/l. Fe(II) and Fe(III) were determined by using the ferrozine method^29^, but instead of reducing Fe(III) with hydroxylamine hydrochloride, ascorbic acid was used ^30^. Absorbance was measured at 562 nm on a spectrophotometer (UV/Vis Spectrometer Lambda 40, Perkin Elmer, Waltham, Massachusetts, USA). The samples were diluted with milli-Q water if needed. The concentration of total P was determined using persulfate digestion^31^. The anion NO_3_^-^ was measured on a Metrohm IC system (883 Basic IC Plus and 919 Autosampler Plus; Riverview, Florida, USA). NO_3_^-^ were separated with a Metrosep A Supp 5 analytical column (250 × 4.0 mm) which was fit with a Metrosep A Supp 4/5 guard column at a flow rate of 0.7 ml/min, using a carbonate eluent (3.2mM Na_2_CO_3_ + 1.0 mM NaHCO_3_). SO_4_ was analyzed using Metrohm IC system (883 Basic IC Plus and 919 Autosampler Plus, Riverview), NH_4_^+^ spectrophotometrically as described by Solórzano^32^, and NO_2_^-^ and DN as in Greenberg et al.^33^.

For the gas analyses, samples from Alaska and Canada were taken as previously described in Kankaala et al.^34^, except that room air was used instead of N_2_ for extracting the gas from the water. Shortly, 30 ml of water was taken into 50 ml syringes, which were warmed to room temperature prior to extraction of the gas. To each syringes 0.5 ml of HNO_3_ and 10 ml of room air was added and the syringes were shaken for 1 minute. Finally, the volumes of liquid and gas phases were recorded and the gas was transferred into glass vials that had been flushed with N_2_ and vacuumed. For Greenland, Sweden and Russia 5 ml of water was taken for the gas samples with a syringe and immediately transferred to 20 ml glass vials filled with N and with 150 μL H_2_PO_4_ to preserve the sample. All gas samples were measured using gas chromatography (Clarus 500, Perkin Elmer, Polyimide Uncoated capillary column 5m × 0.32mm, TCD and FID detector respectively).

### Optical analyses

In order to characterize DOM, we recorded the absorbance of DOM using a UV-visible Carry 100 (Agilent Technologies, Santa Clara, California, USA) or a LAMBDA 40 UV/VIS (PerkinElmer) spectrophotometer, depending on sample origin. SUVA_254_ is a proxy of aromaticity and the relative proportion of terrestrial versus algal carbon sources in DOM^35^ and was determined from DOC normalized absorbance at 254 nm after applying a corrective factor based on iron concentration^36^. S_289_ enlights the importance of fulvic and humic acids related to algal production^37^ and were determined for the intervals 279-299 nm by performing regression calculations using SciLab v 5.5.2.^38^.

We also recorded fluorescence intensity on a Cary Eclipse spectrofluorometer (Agilent Technologies), across the excitation waveband from 250-450 nm (10 nm increments) and emission waveband of 300-560 nm (2 nm increments), or on a SPEX FluoroMax-2 spectrofluorometer (HORIBA, Kyoto, Japan), across the excitation waveband from 250-445 nm (5 nm increments) and emission waveband of 300-600 nm (4 nm increments), depending on sample origin. Based on the fluorometric scans, we constructed excitation-emission matrices (EEMs) after correction for Raman and Raleigh scattering and inner filter effect^39^. We calculated the FI as the ratio of fluorescence emission intensities at 450 nm and 500 nm at the excitation wavelength of 370 nm to investigate the origin of fulvic acids^40^. Higher values (~1.8) indicate microbial derived DOM (autochthonous), whereas lower values (~1.2) indicate terrestrial derived DOM (allochthonous), from plant or soil^41^. HIX is a proxy of the humic content of DOM and was calculated as the sum of intensity under the emission spectra 435–480 nm divided by the peak intensity under the emission spectra 300–445 nm, at an excitation of 250 nm. Higher values of HIX indicate more complex, higher molecular weight, condensed aromatic compounds^42,43^. BIX emphasizes the relative freshness of the bulk DOM. High values (>1) are obtained with the increase of more recently derived DOM, predominantly originated from autochthonous production, while lower values (0.6–0.7) indicate lower production and older DOM^41,43^. BIX was calculated as the ratio of emission at 380 nm divided by the emission intensity maximum observed between 420 and 436 nm at an excitation wavelength of 310 nm^44^.

### High resolution mass spectrometry

50 ml of water samples from each pond was collected and filtered with a Whatman GF/F filter for mass spectrometry analyses. 1.5 ml sample was dried completely with a vacuum drier, and was then re-dissolved in 100 μL 20% acetonitrile, 80% water with three added compounds as internal standards (Hippuric acid, glycyrrhizic acid and capsaicin, all at 400 ppb v/v). Samples were filtered to an autosampler vial and injected at 50 μL onto the column. In order not to overload the detectors, some of the higher concentration samples were injected at a lower volume, to give a maximum of 20μg carbon loaded.

High-performance liquid chromatography – high resolution mass spectrometry (ESI-HRMS) was conducted as described in Patriarca et al.^45^ using a C18-Evo column (100×2.1 mm, 2.6μm; Phenomenex, Torrance, California, USA). The ESI-HRMS data was averaged from 2-17 minutes to allow formula assignment to a single mass list. Formulas considered had masses 150-800 m/z, 4-50 carbon (C) atoms, 4-100 hydrogen (H) atoms, 1-40 oxygen (O) atoms, 0-1 nitrogen (N) atoms and 0-1 13C atoms. Formulas were only considered if they had an even number of electrons, H/C 0.3-2.2 and O/C≤1. The data are presented as number of assigned formulas and weighted average O/C ratio, H/C ratio and m/z. A principal coordinate diagram is also presented, based on pairwise Bray-Curtis dissimilarities of the samples.

On a second run, 24 samples were added from Russia (R1-12) and Greenland (GR1-12). These were treated in the same way as the first 36 samples, and three additional Suwannee River fulvic acid (SRFA, reference material) analyses were performed. At the moment of the run, the DOC concentration of these samples was unknown, so 50μL was injected. From high resolution mass spectrometry, average H/C and number of assigned formulas were obtained. The H/C can be used as a proxy of DOM aliphatic content; higher H/C values (>1) indicate more saturated (aliphatic) compounds, whereas values lower than 1 indicate more unsaturated, aromatic molecules^46^.

### DNA extraction, ITS2 amplification and sequencing

All samples for molecular analyses samples (water and detritus filters and sediments) were extracted with DNeasy PowerSoil^®^ kit (Qiagen, Hilden, Germany), following the manufacturer’s recommendations for low input DNA. Extracts were eluted in 100 μl of Milli-Q water and DNA concentrations were measured with Qubit dsDNA HS kit. The fungal ribosomal internal transcribed spacer 2 (ITS2) sequences were amplified using a modified ITS3 Mix2 forward primer from Tedersoo^47^, named ITS3-mkmix2 CAWCGATGAAGAACGCAG, and a reverse primer ITS4 (equimolar mix of cwmix1 TCCTCCGCTTAyTgATAtGc and cwmix2 TCCTCCGCTTAtTrATAtGc)^14^. Each sample received a unique combination of primers containing identification tags generated by Barcrawl^48^. All tags had a minimum base difference of 3 and a length of 8 nucleotides. Both forward and reverse primer tags were extended by two terminal bases (CA) at the ligation site to avoid bias during ligation of sequencing adaptors, and the forward primer tag also had a linker base (T) added to it^49^. The list of primers and tags is found in Supplementary Table S1. PCR reactions were performed on a final volume of 50 μl, with an input amount of DNA ranging from 0.07 ng to 10 ng, 0.25 μM of each primer, 200 μM of dNTPs, 1U of Phusion™ High-Fidelity DNA Polymerase (Thermo Fisher Scientific, Waltham, Massachusetts, USA), 1X Phusion™ HF Buffer (1X buffer provides 1.5 mM MgCl_2_, Thermo Fisher Scientifics) and 0.015 mg of BSA. PCR conditions consisted of an initial denaturation cycle at 95°C for 3 min, followed by 21-35 cycles for amplification (95°C for 30 sec, 57°C for 30 sec and 72°C for 30 sec), and final extension at 72°C for 10 min. In order to reduce PCR bias, all samples (in duplicates) were first submitted to 21 amplification cycles. In case of insufficient yield, the number of cycles was increased up to 35 cycles (see the records on the number of cycles for each of the samples in Supplementary Table S2).

The PCR products were purified with Sera-Mag™ beads (GE Healthcare Life Sciences, Marlborough, Massachusetts, USA), visualized on a 1.5% agarose gel and quantified using Qubit dsDNA HS kit. The purified PCR products were randomly allocated into three DNA pools (20 ng of each sample), which were purified with E.Z.N.A.^®^ Cycle-Pure kit (Omega Bio-Tek, Norcross, Georgia, USA). Nine of the samples (4 water, 1 sediment and 4 detritus) were left out of the pools because of too little PCR product, giving a total of 203 samples in the pools. Negative PCR controls were added to each pool, as well as a mock community sample containing 10 different fragment sizes from the ITS2 region of a chimera of *Heterobasidium irregular* and *Lophium mytilinum*, ranging from 142 to 591 bases, as described by Castaño et al.^50^. The size distribution and quality of all the pools were verified with BioAnalyzer DNA 7500 (Agilent Technologies), and purity was assessed by spectrophotometry (OD 260:280 and 260:230 ratios) using NanoDrop (Thermo Fisher Scientific). The libraries were sequenced at Science for Life Laboratory (Uppsala University, Sweden), on a Pacific Biosciences Sequel instrument, using 1 SMRT cell per pool.

### Quality filtering of reads, clustering and taxonomy identification of clusters

The sequencing resulted in a total of 1071489 sequences, ranging from 397 to 9184 sequences per sample (average on 2551 sequences per sample). The raw sequences were filtered for quality and clustered using the SCATA pipeline (https://scata.mykopat.slu.se/, accessed on May 19^th^, 2020). For quality filtering, sequences from each pool were screened for the primers and tags, requiring a minimum of 90% match for the primers and a 100% match for the tags. Reads shorter than 100 bp were removed, as well as reads with a mean quality lower than 20, or containing any bases with a quality lower than 7. After this filtering, 281089 sequences were retained in the data. The sequences were clustered at species level by single-linkage clustering at a clustering distance of 1.5%, with penalties of 1 for mismatch, 0 for gap open, 1 for gap extension, and 0 for end gaps. Homopolymers were collapsed to 3 and unique genotypes across all pools were removed. The full UNITE+INSD dataset for Fungi^51^ and customized (provided by Karina E. Clemmensen and Björn D. Lindahl) databases were included in (but without affecting) the clustering process allowing taxonomic affiliation of clusters, hereafter called OTUs (Operational Taxonomic Units), to be determined based on the same criteria as used for the clustering. After the clustering the data included 518128 sequences, divided among 8218 clusters.

## Data Records

The raw sequences are deposited in the NCBI SRA database under accession number PRJNA701021. All the raw mass spectrometry data is available at the Mass Spectrometry Interactive Virtual Environment (MassIVE) under the accession number MSV000086952. The fungal OTUs sequences and their taxonomic classification, as well as all the environmental data related to the samples are deposited in Open Science Framework (OSF) at https://osf.io/hvwpu/?view_only=6f6b73e7522741cc9e350b0d25889db5.

## Technical Validation

The DNA extractions were done in a laminar flow hood with a UV-C lamp and handled separately for each of the locations to avoid any possible cross-contamination between the samples or sites. For the PCRs the samples were first randomized into three groups including samples from all locations to minimize the risk of batch effect at the sequencing step. Negative controls were included in the PCR step and added to the pools. The negative controls created 4, 12 and 4 sequences for pools 1, 2 and 3, respectively. For each pool, 100 ng of a positive control containing mock communities (as described in the methods section) was added. The mock communities captured all different fragment sizes. The sequences were controlled rigorously by removing singletons, and rare reads as described in the methods section to remove any sequences that might be chimeric reads or a result of erroneous amplification. Finally, the sequences were compared against multiple databases and their taxonomy was verified manually to ensure that only fungal sequences were included to the tree shown in Figure 2. For the resulting data, correlations between the number of OTUs and filtered water volume were checked to verify that the differential filtering volumes did not introduce a bias to the richness of the communities (Supplementary Figure S1).

## Usage Notes

To verify that the acquired OTUs were of fungal origin, we scrutinized all the OTUs with a minimum of ten total reads for their phylogenetic origin, retaining 3108 OTUs and 498414 sequences in the analysis. For taxonomic annotation, the Protax-Fungi^52^ and massBLASTer analyses available through the Pluto-F platform of the UNITE database (https://plutof.ut.ee/, accessed on May 23^rd^ 2020) were used. An OTU was assigned to a taxonomic level if the Protax identification probability was at least 95% and matched the taxonomy based on the massBLAST against the UNITE species hypothesis (SH) database. To support the taxonomy identification and discard non-fungal OTUs, a phylogenetic tree was built, which included the OTUs with at least 50 reads in total. For building the tree, the ITS2 region was extracted with ITSx^53^ and aligned with MUSCLE^54^ and a Neighbor-Joining phylogenic tree was constructed using MEGA7^55^, with a p-distance, bootstrap of 1000 replicates, gamma distributed rates and gaps treated as pairwise deletions. As reference sequences, 29 different eukaryotes were selected from the ITS2 database^56^ (http://its2.bioapps.biozentrum.uni-wuerzburg.de/, accessed on August 18^th^ 2020 – accession numbers: AB084092, AY752993, HQ219352, GQ402831, AF228083, AF053158, AF163102, JN113133, AF353997, GU001158, JQ340345, AM396560, FJ946912, EU812490, KJ925151, KF772413, JX988759, AF315074, AJ400496, AJ566147, AY458037, AY479922, EF060369, FN397599, JF750409, KF524372, U65485, Z48468, AY499004). Eight references were further derived from GenBank (https://www.ncbi.nlm.nih.gov/genbank/, accessed on September 29^th^ 2020 - accession numbers: KC357673, AF508774, AY676020, MN158348, HM161704, AY264773, AB906385, AJ296818). These sequences cover different eukaryote lineages: Centroheliozoa, Choanoflagellida, Ichthyosporea, Oomycetes, Streptophyta, Chlorophyta, Rhodophyta, Cercozoa, Amoebozoa, Apusozoa, Cryptophyta, Haptophyceae, Heterolobosea, Katablepharidophyta, Arthropoda, Picozoa, Alveolata, Cnidaria, Stramenopiles, Protostomia and Porifera. Additionally, 530 fungal sequences (SHs from UNITE database) were selected to represent different fungal lineages and included the closest SH matches to each of the OTUs. Finally, to further establish higher level taxonomy, BLASTn searches (E-value cutoff of 1e-3, against the NCBI nucleotide database (https://www.uppmax.uu.se/resources/databases/blast-databases/), were run on September-October 2020 excluding uncultured organisms and environmental samples) for all the OTUs. The (at least) 10 best hits from the BLASTn search were evaluated as follows: an OTU was classified as “likely fungi” if all resulting hits would match fungal sequences. The OTUs that have also matched other eukaryotic sequences were checked manually, and classified as “likely fungi” if the best hits were fungal sequences, and also based on the quality of the alignments (higher query coverage and identity %). Any OTU that could not be classified as “likely fungi” from the BLASTn searches and/or would cluster with any other eukaryote reference in the phylogenetic tree was discarded. The final taxonomic assignment of all fungal OTUs was based on matches to UNITE SHs and Protax probabilities. OTUs with undetermined phylum, class or order were assigned to higher taxonomic levels whenever supported by the phylogenetic tree (considering a minimal bootstrap value of 70%) or BLASTn results (min of 90% query coverage and 90% identity for class, and 100% query coverage and min 97% identity for order). The final data set had 1334 OTUs (178531 sequences) identified as fungi and are presented in Supplementary Figure 2.

## Code Availability

The code used for the extraction of ITS2 regions using ITSx and for the Blastn searches are provided in Supplementary Note 1.

## Acknowledgements

We thank the staff at the Toolik Field Station, Arctic Station, CEN W-K Station, Abisko Research Station and Khanymey Station, as well as Anelise Kluge, Gaëtan Martin and Martin Petersson for their assistance with the samplings. We offer our gratitude to the Cree and Inuit communities in Whapmagoostui-Kuujjuarapik for giving us access to their ancestral lands. Katarina Ihrmark, Maria Jonsson, Christoffer Bergvall and Anna Ferguson are acknowledged for their assistance with the laboratory analyses. We are grateful to the INTERACT and Science for Life Laboratory for funding. The computations were performed on resources provided by SNIC through Uppsala Multidisciplinary Center for Advanced Computational Science (UPPMAX) under Project SNIC 2020-5-196.

## Author contributions

MK planned the samplings together with CW and SP, did the molecular laboratory work and sequence analyses and wrote the first draft of the manuscript together with SP CW planned the study together with SP, planned the sampling together with MK and SP, participated in the collection of the samples

MW participated in the collection of the samples, analyzed carbon samples and data

KEC instructed the amplicon processing

JH analyzed carbon samples and data

KE analyzed carbon samples and data

JS helped with the study design and data analysis

SP planned the study together with CW, planned the samplings together with CW and MK, participated in the collection of the samples and wrote the first draft of the manuscript together with MK

All authors participated in the revision of the manuscript

## Competing interests

No competing interests.

## Figure Legends

Figure 1. Locations of the sampling sites.

Figure 2. Workflow for producing the amplicon and chemical data.

## Table Legends

Table 1. Description of the sampled sites. The stations are ordered according to increasing level of thermokarst activity. The coordinates are for the stations, the sample coordinates can be found at https://osf.io/hvwpu/?view_only=6f6b73e7522741cc9e350b0d25889db5.

## Supplementary Materials

Supplementary Figure S1. Filtered volumes plotted against the number of OTUs.

Supplementary Figure S2. Neighbor-Joining phylogenetic tree of the fungal OTUs and fungal references sequences obtained from UNITE database.

Supplementary Table S1. The barcodes used for creating the sequencing libraries.

Supplementary Table S2. The number of PCR cycles used to create the sequencing libraries.

Supplementary Note 1. Code for extracting ITS2 regions with ITSx and Blastn searches.

Supplementary Figure S1. Filtered volumes plotted against the number of OTUs. Sampled volume (ml) x number of observed OTUs, per sample. Red dots are samples filtered with 0.22 μm filters, and blue dots represent 5 μm filters. Spearman correlations between sample volume and number of observed OTUs was 0.30 for 5 μm filtered samples, and 0.01 for 0.22 μl filters.

Supplementary Figure S2. Neighbor-Joining phylogenetic tree of the fungal OTUs and fungal references sequences obtained from UNITE database. In the figure, Ascomycetes are shown in blue, Basidiomyces in red, Chytridiomycetes in orange and Mucoromycetes in pink.

## Notes

https://osf.io/hvwpu/?view_only=6f6b73e7522741cc9e350b0d25889db5

### Competing Interest Statement

The authors have declared no competing interest.

