## Supplementary figures and images for "Fungal community composition along a gradient of permafrost thaw"

### Supplementary Figure 2

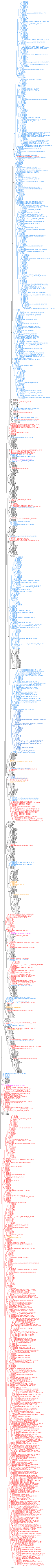
