## Supplementary Table 2 for "Fungal community composition along a gradient of permafrost thaw"

**Pool 1** 1

| | | | | | | | | | | | | | | | | | | | | | | |
|---|---|---|---|---|---|---|---|---|---|---|---|---|---|---|---|---|---|---|---|---|---|---|
| A - 1 | **T1 0.2 31c** | A - 2 | **SAS2C 5 28c** | A - 3 | **R10 0.2 28c** | A - 4 | **GR3 5 28c** | A - 5 | **P1 sed 21c** | A - 6 | | A - 7 | **T2 d 28c** | A - 8 | **T9 sed 22c** | A - 9 | **GR3 sed 21c** | A - 10 | **P10 5 20c** | A - 11 | **P2 d 22c** | A - 12 |
| B - 1 | **T3 0.2 28c** | B - 2 | **B4 5 28c** | B - 3 | **R3 5 28c** | B - 4 | **GR6 5 28c** | B - 5 | **P2 sed 20c** | B - 6 | | B - 7 | **T4 d 28c** | B - 8 | **T10 sed 22c** | B - 9 | **GR4 sed 21c** | B - 10 | **P11 5 21c** | B - 11 | **P4 d 25c** | B - 12 |
| C - 1 | **T4 0.2 28c** | C - 2 | **P9 0.2 28c** | C - 3 | **R7 5 23c** | C - 4 | **GR8 5 28c** | C - 5 | **P3 sed 23c** | C - 6 | | C - 7 | **T7 d 28c** | C - 8 | **T11 sed 25c** | C - 9 | **GR5 sed 23c** | C - 10 | **6 sed 21c** | C - 11 | **P5 d 23c** | C - 12 |
| D - 1 | **T9 0.2 35c** | D - 2 | **P9 5 29c** | D - 3 | **GR1 0.2 28c** | D - 4 | **P1 0.2 24c** | D - 5 | **P9 sed 23c** | D - 6 | | D - 7 | **T8 d 28c** | D - 8 | **T12 sed 25c** | D - 9 | **GR6 sed 23c** | D - 10 | **9 pond C2 sed 22c** | D - 11 | **P6 d 25c** | D - 12 |
| E - 1 | **T1 5 31c** | E - 2 | **P12 5 28c** | E - 3 | **GR2 0.2 28c** | E - 4 | **P4 0.2 27c** | E - 5 | **R3 sed 20c** | E - 6 | | E - 7 | **GR1 d 28c** | E - 8 | **B4 0.2 29c** | E - 9 | **I1 5 25c** | E - 10 | **10 pond B1 sed 22c** | E - 11 | | E - 12 |
| F - 1 | **T2 5 28c** | F - 2 | **R8 0.2 8c** | F - 3 | **GR3 0.2 28c** | F - 4 | **P8 0.2 27c** | F - 5 | **R5 sed 20c** | F - 6 | | F - 7 | **GR2 d 28c** | F - 8 | **C2 0.2 27c** | F - 9 | **I3 5 25c** | F - 10 | **11 sed 22c** | F - 11 | | F - 12 |
| G - 1 | **T3 5 27c** | G - 2 | **R4 0.2 28c** | G - 3 | **GR4 0.2 28c** | G - 4 | **R4 5 25c** | G - 5 | **R6 sed 23c** | G - 6 | | G - 7 | **GR5 d 28c** | G - 8 | **C5 0.2 29c** | G - 9 | | G - 10 | neg | G - 11 | | G - 12 |
| H - 1 | **T5 5 24c** | H - 2 | **R7 0.2 28c** | H - 3 | **GR2 5 31c** | H - 4 | **R2 5 24c** | H - 5 | **R7 sed 23c** | H - 6 | | H - 7 | **GR6 d 28c** | H - 8 | **I1 0.2 25c** | H - 9 | | H - 10 | | H - 11 | | H - 12 |

**Pool 2**

| | | | | | | | | | | | | | | | | | | | | | | |
|---|---|---|---|---|---|---|---|---|---|---|---|---|---|---|---|---|---|---|---|---|---|---|
| A - 1 | **T2 0.2 28c** | A - 2 | **T4 sed 27c** | A - 3 | **SAS2A 5 26c** | A - 4 | **P11 0.2 28c** | A - 5 | **R1 5 26c** | A - 6 | **12 pond G1 sed 20c** | A - 7 | **GR9 5 30c** | A - 8 | **P7 5 24c** | A - 9 | | A - 10 | **T3 d 31c** | A - 11 | **T1 d 31c** | A - 12 |
| B - 1 | **T5 0.2 28c** | B - 2 | **T5 sed 25c** | B - 3 | **SAS2B 5 25c** | B - 4 | **P1 5 25c** | B - 5 | **R6 5 21c** | B - 6 | **14 sed 21c** | B - 7 | **GR10 5 30c** | B - 8 | **R10 sed 25c** | B - 9 | | B - 10 | **T5 d 28c** | B - 11 | **GR3 d 31c** | B - 12 |
| C - 1 | **T6 0.2 28c** | C - 2 | **T7 sed 27c** | C - 3 | **SAS2D 5 25c** | C - 4 | **P3 5 25c** | C - 5 | **R8 5 22c** | C - 6 | **19 sed 22c** | C - 7 | **GR11 5 31c** | C - 8 | | C - 9 | neg | C - 10 | **T6 d 25c** | C - 11 | | C - 12 |
| D - 1 | **T7 0.2 31c** | D - 2 | **T8 sed 27c** | D - 3 | **G1 5 28c** | D - 4 | **P8 5 27c** | D - 5 | **R11 5 25c** | D - 6 | **P10 sed 21c** | D - 7 | **GR12 5 32c** | D - 8 | | D - 9 | | D - 10 | **P1 d 25c** | D - 11 | | D - 12 |
| E - 1 | **T4 5 25c** | E - 2 | **SAS2A 0.2 31c** | E - 3 | **1 sed 28c** | E - 4 | **R1 0.2 28c** | E - 5 | GR11 0.2 Failed* | E - 6 | **P11 sed 22c** | E - 7 | **GR12 sed 21c** | E - 8 | | E - 9 | | E - 10 | **P3 d 25c** | E - 11 | | E - 12 |
| F - 1 | **T7 5 29c** | F - 2 | **SAS2B 0.2 28c** | F - 3 | **P2 0.2 27c** | F - 4 | **R3 0.2 25c** | F - 5 | **GR9 0.2 31c** | F - 6 | **R8 sed 21c** | F - 7 | **GR7 sed 22c** | F - 8 | | F - 9 | | F - 10 | **P7 d 26c** | F - 11 | | F - 12 |
| G - 1 | **T8 5 28c** | G - 2 | **SAS2C 0.2 31c** | G - 3 | **P3 0.2 29c** | G - 4 | **R9 0.2 25c** | G - 5 | **GR10 0.2 28c** | G - 6 | R11 sed Failed | G - 7 | **GR8 sed 22c** | G - 8 | | G - 9 | **P12 d 25c** | G - 10 | GR4 d Failed | G - 11 | | G - 12 |
| H - 1 | **P4 sed 28c** | H - 2 | **SAS2D 0.2 31c** | H - 3 | **P5 0.2 28c** | H - 4 | **R2 0.2 29c** | H - 5 | GR12 0.2 Failed | H - 6 | **R12 sed 21c** | H - 7 | **GR9 sed 22c** | H - 8 | | H - 9 | **P12 sed 25c** | H - 10 | GR10 d Failed | H - 11 | **GR12 d 30c** | H - 12 |

**Pool 3**

| | | | | | | | | | | | | | | | | | | | | | | | |
|---|---|---|---|---|---|---|---|---|---|---|---|---|---|---|---|---|---|---|---|---|---|---|---|
| A - 1 | **T6 5 22c** | A - 2 | **C5 5 23c** | A - 3 | **P5 5 26c** | A - 4 | **R5 5 22c** | A - 5 | **GR11 sed 22c** | A - 6 | **T10 0.2 28c** | A - 7 | **P2 5 20c** | A - 8 | | A - 9 | **P12 0.2 31c** | A - 10 | | A - 11 | **T9 d 28c** | A - 12 | **GR7 d 30c** |
| B - 1 | **T1 sed 25c** | B - 2 | **H1 5 232c** | B - 3 | **P6 5 23c** | B - 4 | **R12 5 22c** | B - 5 | **GR5 0.2 28c** | B - 6 | **T11 0.2 28c** | B - 7 | **P4 5 22c** | B - 8 | | B - 9 | T8 0.2 Failed | B - 10 | | B - 11 | **T10 d 28c** | B - 12 | GR8 d Failed |
| C - 1 | **T2 sed 22c** | C - 2 | **8 pond C5 sed 23c** | C - 3 | **P5 sed 25c** | C - 4 | **R1 sed 25c** | C - 5 | **GR6 0.2 28c** | C - 6 | **T12 0.2 31c** | C - 7 | **P10 0.2 22c** | C - 8 | **G1 0.2 27c** | C - 9 | neg | C - 10 | | C - 11 | **T11 d 28c** | C - 12 | **GR9 d 28c** |
| D - 1 | **T3 sed 25c** | D - 2 | **16 sed 25c** | D - 3 | **P6 sed 28c** | D - 4 | **R2 sed 25c** | D - 5 | **GR7 0.2 25c** | D - 6 | **T9 5 28c** | D - 7 | **GR10 sed 20c** | D - 8 | **H1 0.2 25c** | D - 9 | | D - 10 | | D - 11 | | D - 12 | GR11 d Failed |
| E - 1 | **T6 sed 26c** | E - 2 | **17 sed 25c** | E - 3 | **P7 sed 25c** | E - 4 | **R4 sed 25c** | E - 5 | **GR8 0.2 25c** | E - 6 | **T10 5 28c** | E - 7 | **GR1 5 31c** | E - 8 | | E - 9 | | E - 10 | | E - 11 | **P8 d 22c** | E - 12 | |
| F - 1 | **B1 0.2 26c** | F - 2 | **21 "tiny" sed 25c** | F - 3 | **P8 sed 25c** | F - 4 | **R9 sed 25c** | F - 5 | **R5 0.2 25c** | F - 6 | **T11 5 25c** | F - 7 | **GR4 5 32c** | F - 8 | | F - 9 | | F - 10 | | F - 11 | **P9 d 24c** | F - 12 | |
| G - 1 | **B1 5 23c** | G - 2 | **P6 0.2 27c** | G - 3 | **R9 5 29c** | G - 4 | **GR1 sed 25c** | G - 5 | **R6 0.2 28c** | G - 6 | R11 0.2 Failed | G - 7 | **GR5 5 31c** | G - 8 | | G - 9 | | G - 10 | | G - 11 | **P10 d 24c** | G - 12 | |
| H - 1 | **C2 5 23c** | H - 2 | **P7 0.2 27c** | H - 3 | **R10 5 22c** | H - 4 | **GR2 sed 25c** | H - 5 | **I3 0.2 27c** | H - 6 | **R12 0.2 29c** | H - 7 | **GR7 5 32c** | H - 8 | | H - 9 | | H - 10 | | H - 11 | **P11 d 24c** | H - 12 | |

*samples that did not amplify and were left out of the pools for sequencing
