## Supplementary Figure 1, Supplementary Table 1, Supplementary Note 1 for "Fungal community composition along a gradient of permafrost thaw"

**Supplementary Information**

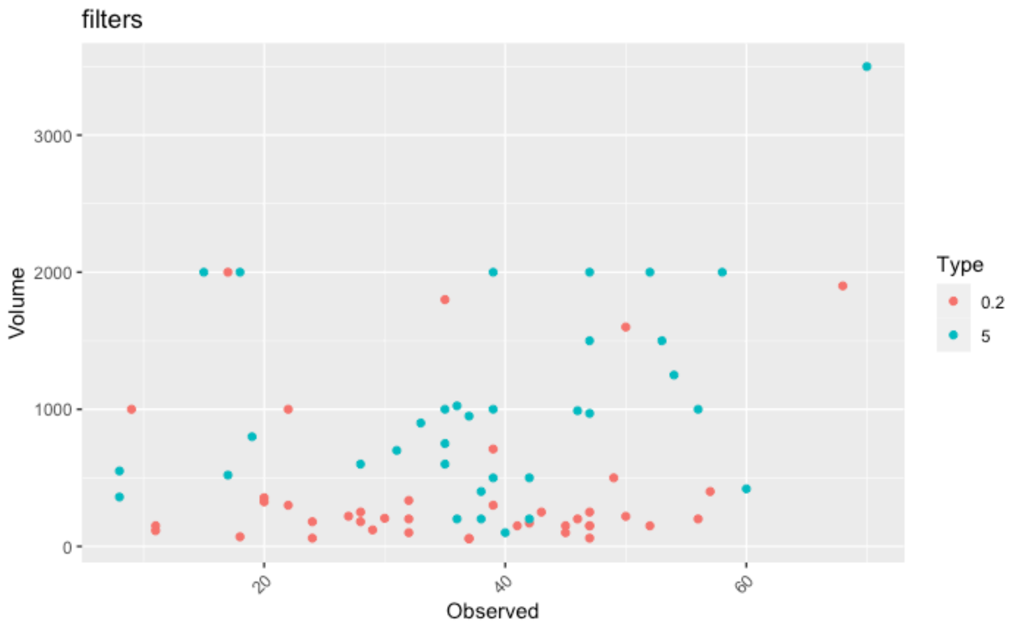


**Supplementary Figure S1** – Filtered volumes plotted against the number of OTUs. Sampled volume (ml) x number of observed OTUs, per sample. Red dots are samples filtered with 0.22 µm filters, and blue dots represent 5 µm filters. Spearman correlations between sample volume and number of observed OTUs was 0.30 for 5 µm filtered samples, and 0.01 for 0.22 µl filters.

**Supplementary Table S1** – The barcodes used for creating the sequencing libraries.

| Primer_ID with tag name | Tag **(bold)** and primer sequences |
|---|---|
| Forward tagged primers: | |
| ITS3-mkmix2-A | **CATACCAGCAT**CAWCGATGAAGAACGCAG |
| ITS3-mkmix2-B | **CATCACGACAT**CAWCGATGAAGAACGCAG |
| ITS3-mkmix2-C | **CAAAGACCGAT**CAWCGATGAAGAACGCAG |
| ITS3-mkmix2-D | **CACAACACCAT**CAWCGATGAAGAACGCAG |
| ITS3-mkmix2-E | **CACTAGCACAT**CAWCGATGAAGAACGCAG |
| ITS3-mkmix2-F | **CACATGTCCAT**CAWCGATGAAGAACGCAG |
| ITS3-mkmix2-G | **CAGGATCACAT**CAWCGATGAAGAACGCAG |
| ITS3-mkmix2-H | **CAGGCAAGTAT**CAWCGATGAAGAACGCAG |
| Reverse tagged primers: | |
| ITS4-cwmix1-01 | **CAAACCTCCA**TCCTCCGCTTAY*TG*ATAT*GC |
| ITS4-cwmix2-01 | **CAAACCTCCA**TCCTCCGCTTAT*TR*ATAT*GC |
| ITS4-cwmix1-02 | **CATGAACGAC**TCCTCCGCTTAY*TG*ATAT*GC |
| ITS4-cwmix2-02 | **CATGAACGAC**TCCTCCGCTTAT*TR*ATAT*GC |
| ITS4-cwmix1-03 | **CATCTCAGAC**TCCTCCGCTTAY*TG*ATAT*GC |
| ITS4-cwmix2-03 | **CATCTCAGAC**TCCTCCGCTTAT*TR*ATAT*GC |

| | |
|---|---|
| ITS4-cwmix1-04 | **CAATCTAGCC**TCCTCCGCTTAY*TG*ATAT*GC |
| ITS4-cwmix2-04 | **CAATCTAGCC**TCCTCCGCTTAT*TR*ATAT*GC |
| ITS4-cwmix1-05 | **CAACTAGGAC**TCCTCCGCTTAY*TG*ATAT*GC |
| ITS4-cwmix2-05 | **CAACTAGGAC**TCCTCCGCTTAT*TR*ATAT*GC |
| ITS4-cwmix1-06 | **CAAGCAGTGA**TCCTCCGCTTAY*TG*ATAT*GC |
| ITS4-cwmix2-06 | **CAAGCAGTGA**TCCTCCGCTTAT*TR*ATAT*GC |
| ITS4-cwmix1-07 | **CACTGAGGAA**TCCTCCGCTTAY*TG*ATAT*GC |
| ITS4-cwmix2-07 | **CACTGAGGAA**TCCTCCGCTTAT*TR*ATAT*GC |
| ITS4-cwmix1-08 | **CACAATGGTG**TCCTCCGCTTAY*TG*ATAT*GC |
| ITS4-cwmix2-08 | **CACAATGGTG**TCCTCCGCTTAT*TR*ATAT*GC |
| ITS4-cwmix1-09 | **CACGATGAGA**TCCTCCGCTTAY*TG*ATAT*GC |
| ITS4-cwmix2-09 | **CACGATGAGA**TCCTCCGCTTAT*TR*ATAT*GC |
| ITS4-cwmix1-10 | **CACGAACAAG**TCCTCCGCTTAY*TG*ATAT*GC |
| ITS4-cwmix2-10 | **CACGAACAAG**TCCTCCGCTTAT*TR*ATAT*GC |
| ITS4-cwmix1-11 | **CACGCACTAA**TCCTCCGCTTAY*TG*ATAT*GC |
| ITS4-cwmix2-11 | **CACGCACTAA**TCCTCCGCTTAT*TR*ATAT*GC |
| ITS4-cwmix1-12 | **CAGCTATGAG**TCCTCCGCTTAY*TG*ATAT*GC |
| ITS4-cwmix2-12 | **CAGCTATGAG**TCCTCCGCTTAT*TR*ATAT*GC |

Bases followed by * are PTO-protected nucleotides that prevent mismatch corrections by proof-reading polymerases.

**Supplementary Note 1 –** Code for extracting ITS2 regions with ITSx and Blastn searches.

ITSx command line:

```
ITSx -i 3108_OTUs_scata4743.fasta -o ITSx_3108_OTUs_scata4743 --save_regions all
```

Blastn command line:

```
blastn -db nt -query 3108_OTUs_scata4743.fasta -negative_gilist sequence.gi -evalue 1e-3 -max_target_seqs 10 -max_hsps 10 -out Result_blastn_OTUs_scata4743 -outfmt "6 qseqid sseqid pident length mismatch gapopen qstart qend sstart send evalue bitscore staxids sscinames scomnames sblastnames sskingdoms stitle qcovs"
```
